# sPepFinder expedites genome-wide identification of small proteins in bacteria

**DOI:** 10.1101/2020.05.05.079178

**Authors:** Lei Li, Yanjie Chao

## Abstract

Small proteins shorter than 50 amino acids have been long overlooked. A number of small proteins have been identified in several model bacteria using experimental approaches and assigned important functions in diverse cellular processes. The recent development of ribosome profiling technologies has allowed a genome-wide identification of small proteins and small ORFs (smORFs), but our incomplete understanding of small proteins hinders *de novo* computational prediction of smORFs in non-model bacterial species. Here, we have identified several sequence features for smORFs by a systematic analysis of all the known small proteins in *E. coli*, among which the translation initiation rate is the strongest determinant. By integrating these features into a support vector machine learning model, we have developed a novel sPepFinder algorithm that can predict conserved smORFs in bacterial genomes with a high accuracy of 92.8%. *De novo* prediction in *E. coli* has revealed several novel smORFs with evidence of translation supported by ribosome profiling. Further application of sPepFinder in 549 bacterial species has led to the identification of > 100,000 novel smORFs, many of which are conserved at the amino acid and nucleotide levels under purifying selection. Overall, we have established sPepFinder as a valuable tool to identify novel smORFs in both model and non-model bacterial organisms, and provided a large resource of small proteins for functional characterizations.

## INTRODUCTION

Small proteins less than 50 amino acids have been increasingly appreciated to play critical roles in diverse biological processes in all three kingdoms of life (Andrews and Rothnagel, 2014). Despite lacking annotations in bacterial genomes, small proteins encoded by open reading frames (smORFs) have been fortuitously identified in the gaps between protein-coding genes, sometime even inside small noncoding RNAs (Jørgensen et al., 2013; Pinel-Marie et al., 2014). Most of the known bacterial small proteins are leader peptides upstream of a structural gene, or interaction partners of larger protein (complexes), or small toxic proteins as part of the toxin/antitoxin system (TA system) (Hemm et al., 2008). The functions of those bacterial small proteins have been assigned in a myriad of physiological pathways including biofilm formation, stress response, antibiotic resistance, virulence and beyond (Duval and Cossart, 2017; Storz et al., 2014).

The identification of smORFs has been challenging due to their small size and a lack of sequence/structural characteristics. Traditionally, bacterial small proteins were identified individually by biochemical isolation (Levin et al., 1993). With increasing availability of complete genome sequences, smORFs have been considered as ‘noise’ and intentionally left out during genomic annotation of ORFs by automatic computational programs. Very few tools were able to accurately predict smORFs directly from genome sequence (Baek et al., 2017; Boekhorst et al., 2011; Pauli et al., 2015; Zhu and Gribskov, 2019). It was only recently that a combination of computational screening and experimental approaches identified an array of small proteins in the model bacterium E. coli (Hemm et al., 2008) (Miravet-Verde et al., 2019).

The recently developed ribosomal profiling approach (a.k.a. Ribo-seq of ribosome-protected mRNA fragments) has become instrumental to detect novel ORFs in both eukaryotic and prokaryotic organisms (Brar and Weissman, 2015; Ingolia, 2014; Weaver et al., 2019). Ribo-seq data could provide evidence of active translation, enabling straight-forward predictions of novel ORFs as well as smORFs with high confidence (Aspden et al., 2014; Bazzini et al., 2014; Calviello et al., 2016; Ji et al., 2015; Raj et al., 2016). However, such experimental approach is labour-intensive and restricted to a few model bacterial species such as *E. coli* and *Bacillus subtilis* (Li et al., 2012; Oh et al., 2011; Sberro et al., 2019; Schrader et al., 2014; Woolstenhulme et al., 2015). The low resolution of Ribo-seq analysis in bacteria complicates the precise determination of reading frames and starting codons. The resolution could be improved using some modified protocols (Weaver et al., 2019), which however inevitably introduce additional biases and costs. Therefore, other alternative approaches are in need to allow the rapid identification of smORFs in a wide range of bacteria.

Here, we have identified three strong features of smORF that allow the discrimination of smORFs from random ORFs in bacterial chromosomes. We have developed upon these features a support vector machine learning model (sPepfinder) to predict smORF with 92.8% accuracy. sPepfinder can be used as a stand-alone tool for the prediction of smORFs from genome sequences with high-confidence, or used in combination with ribosome profiling to accurately predict translating smORFs. The development of sPepfinder promises a high-throughput identification of smORFs in diverse bacterial species for future functional studies.

## RESULTS

### Systematic analyses revealed distinct features for smORF

Most small proteins lack functional domains and remain uncharacterized. To better understand the features of small proteins, we sought to systematically analyse 94 known small proteins in *Escherichia coli* from EcoGene database (version: 3.0) (Zhou and Rudd, 2012). These small proteins can be classified into several categories based on genetic locations. Most of the small proteins are encoded by independent genes in the intergenic regions. Less than a quarter partially overlap with other genes or pseudogenes, and only 7% are leader peptides linked with the gene downstream. A vast majority of known small proteins (96%) use a canonical start codon ATG or GTG, with an average length of 30 amino acids (Figure 1A). Interestingly, our analysis of amino acid frequency revealed that three amino acids are significantly enriched in small proteins (student *t* test: *p*-value < 0.001, Figure 1B). All three amino acids (Leucine, Isoleucine and Valine) are hydrophobic amino acids, in line with previous observations that many small proteins are localized in the membrane (Alix and Blanc-Potard, 2009; Storz et al., 2014).

**Figure 1.**
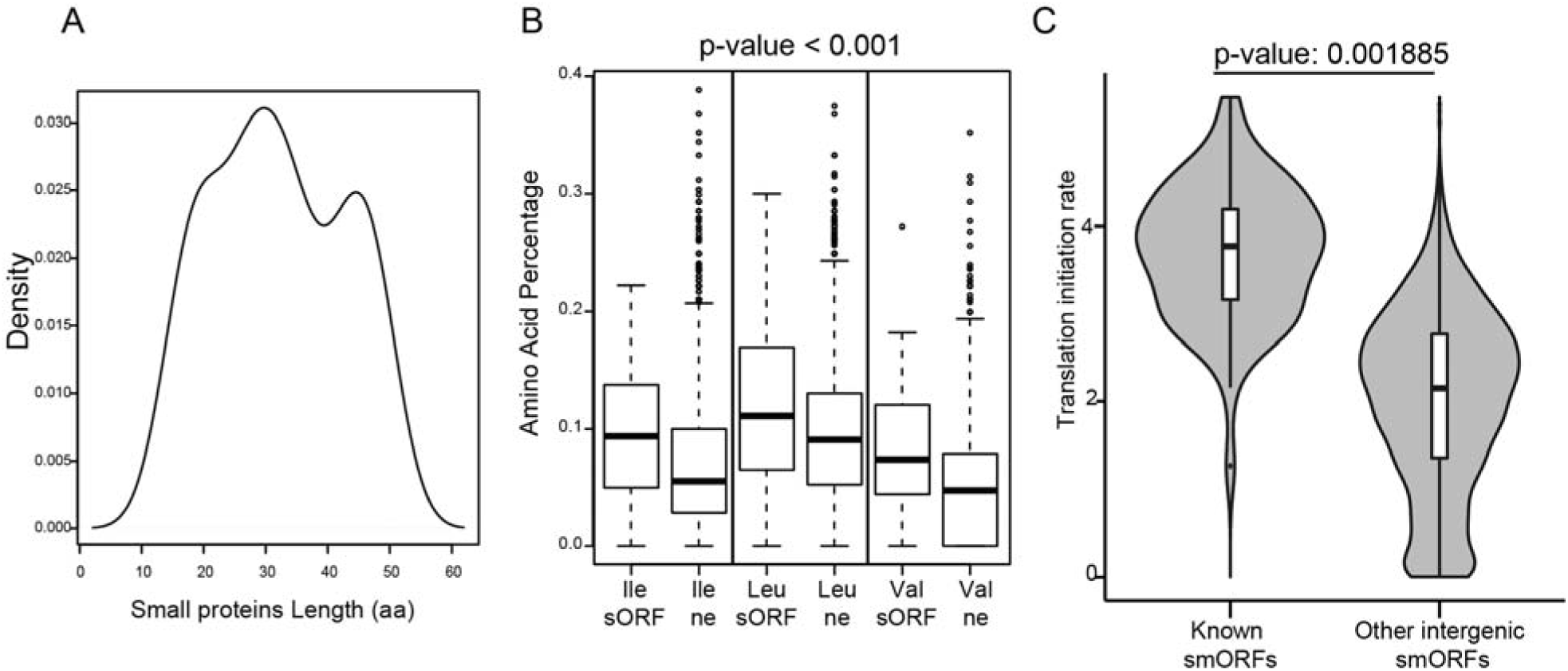
Bacterial smORFs have distinct sequence features. (A) Size distribution of known small proteins in *Escherichia coli*. (B) The percentage of hydrophobic amino acids in small proteins. The negative control (NE) are random short ORFs in the chromosome. (C) The translation initiation score for known small proteins and random intergenic short ORFs in *E. coli*.

The ribosome binding site (RBS) is the key sequence element that determines the translation efficiency of proteins. To understand the translation efficiency of small proteins in *E. coli*, we have analysed their RBS strength and translation initiation rate using a thermodynamic model, which evaluates the Gibbs free energy of molecular interactions between RBS and the 16S ribosomal RNA (Salis et al., 2009). Strikingly, we found that most small proteins possess stronger RBS with significantly higher translation initiation scores, compared to that of random ORFs predicted in the intergenic regions (student *t* test: *p*-value: 0.001885, Figure 1C). In fact, the top quartile of known small proteins has stronger RBS than most, if not all, the intergenic short ORFs. Further analysis showed that weaker RBS (bottom quartile) are often associated with small toxin proteins and anti-toxins (TA systems), including ldrA, ldrB, ldrC, whose functions may require lower expression (Fozo *et al*, 2008).

### sPepfinder employs a machine learning model to predict smORFs

Having identified these features for small proteins, we next established a machine learning algorithm to model on these features and predict new smORFs (Figure 2A). Specifically, a support vector model was built by training on a positive dataset that contains 110 non-redundant known small proteins collected from ten different Gamma-proteobacterial species including *E. coli*, and a negative dataset that contains 136 short ORF predicted in rRNAs and tRNAs in these representative strains (Supplemental Table S1). The input parameters included the RBS strength (the translation initiation score), the frequency of hydrophobic amino acids (I, L, V), and their coding potential scores based on the sequence conservation of codons (RNAcode, (Washietl et al., 2011).

**Figure 2.**
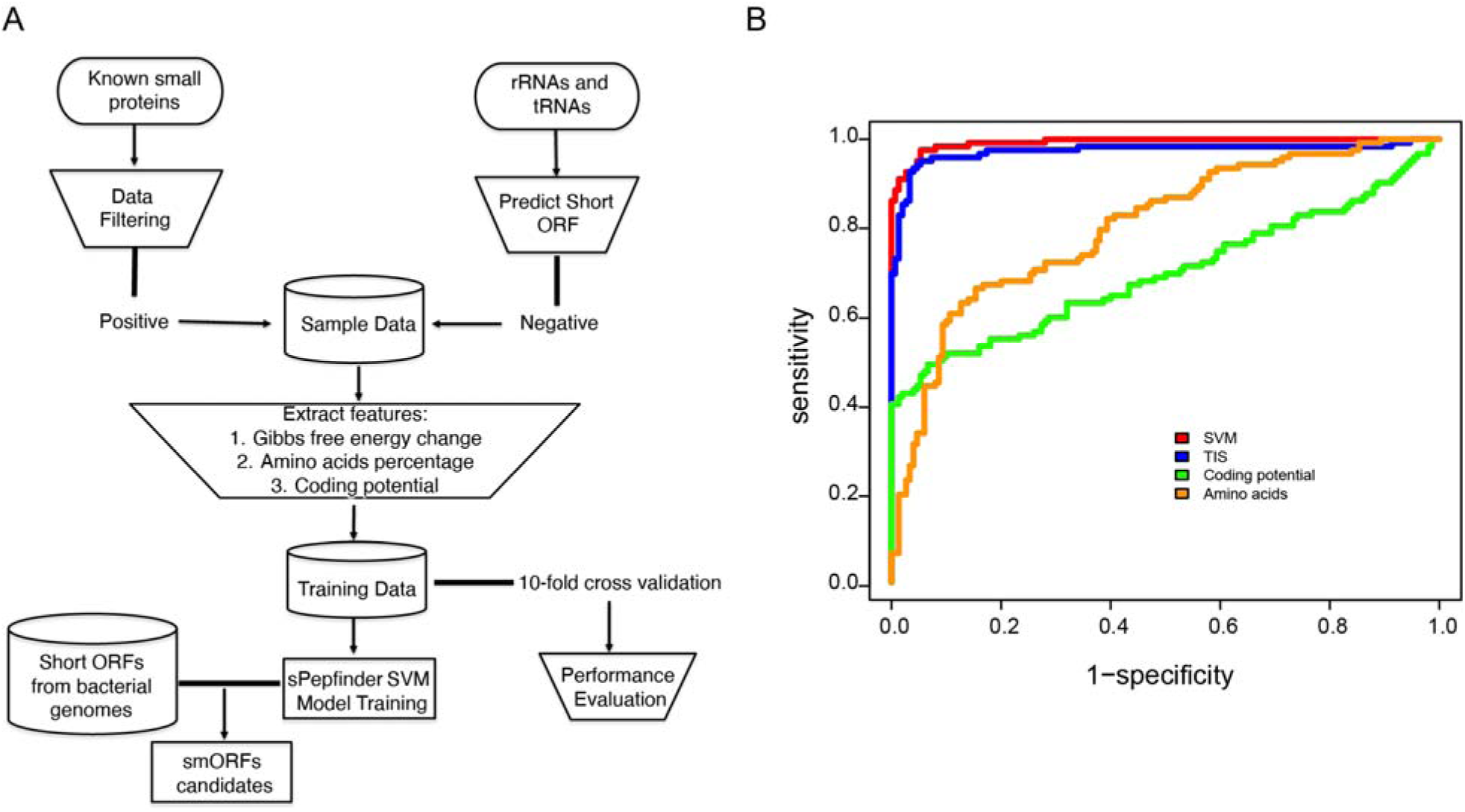
The identification of bacterial smORFs using sPepfinder. (A) Schematic framework of the sPepfinder algorithm. (B) ROC evaluation of sPepfinder performance with different features, including TIS (transcription initiation score), coding conservation potential and hydrophobic amino acid frequency.

In order to evaluate the performance of our algorithm, we have carried out a ROC (Receiver operating characteristic) analysis with 10-fold cross-validation (Figure 2B). The training sets were split into ten datasets, and then 90% of them was used a training model and the remaining 10% as test datasets. The ROC results show that our SVM model reached 92.8% accuracy on predicting the true-positives and the true-negatives. Among all three features of small proteins, the translation initiation score is the most predominant SVM classifier (Figure 2B). Thus, the RBS strength is the strongest indicator of prediction confidence, whereas the additional sequence features help define the frame of translation and the size of small proteins.

### *De novo* prediction of smORFs in *Escherichia coli*

To further validate our approach, we have used sPepfinder to predict smORFs in the genome of *Escherichia coli* MG1655, and evaluated the expression of small proteins using the available ribosome profiling data in the GWIPS-viz database (Michel et al., 2014). In total, sPepfinder has found 124 smORFs using a SVM score cutoff at probability of 0.9 (Supplemental Table S2). Interestingly, more than 40% of them are known small proteins (Figure 3A), including several known small proteins that were not included in our initial training set such as YohO, a known membrane-bound small protein (Hemm et al., 2008). sPepfinder has successfully predicted the correct size and the reading frame of YohO (Figure 3B), highlighting the accuracy of our SVM model.

**Figure 3.**
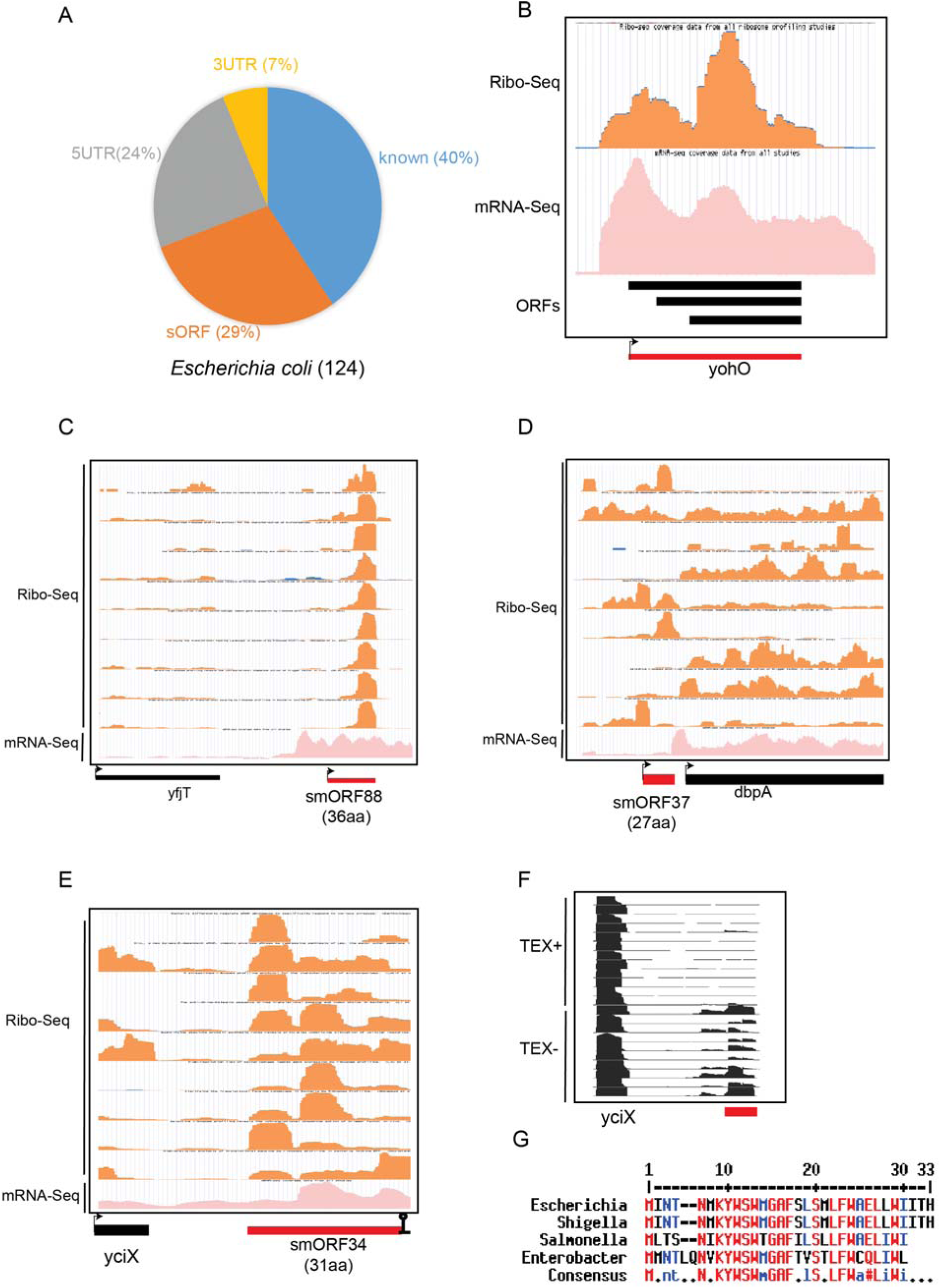
sPepfinder identifies 104 smORFs in *Escherichia coli*. (A) Pie chart distribution of 124 predicted smORFs in *Escherichia coli*. (B) The Ribo-seq and mRNA-seq coverage profiles for the known small protein YohO, shown as a red bar. Black bars indicate alternative frames of translation. (C-E) The Ribo-seq and mRNA-seq coverage profiles for new small proteins smORF88 (C), smORF37 (D), and smORF34 (E). (F) differential RNA-seq profiles for the transcripts of smORF34. TEX+/− indicates the treatment of 5’p-dependent terminator exoribonuclease. (G)The sequence conservation of smORF34 among the close-related enterobacterial species.

sPepfinder has not only identified novel smORFs in the intergenic regions (29%), but also novel smORFs in the 5’UTRs and 3’UTRs (Figure 3A), many of which are supported by ample amount of ribosome footprints as well as RNA-seq reads at different conditions. These include an smORF downstream of *yfjT* (Figure 3C), a putative uORF (Fig. 3D) in the 5’UTR of *dbpA* encoding an ATP-dependent RNA helicase (Diges & Uhlenbeck, 2005), as well as an smORF in the 3’UTR of *yciX* (Fig. 3E). The latter 31aa-smORF is conserved across enterobacteria (Figure 3G), and is encoded on the highly expressed *yciX* 3’UTR without an independent promoter (Figure 3F), according to the available differential RNA-seq data (Thomason et al., 2015).

### Genome-wide prediction of smORFs in diverse bacteria

The successful application of sPepfinder in *E. coli* has promoted us to predict smORF in other related bacterial species. Using the same strict cutoff (probability > 0.9), we have harnessed sPepfinder to identify smORFs in 549 bacterial genomes representing the full spectrum of Proteobacteria phylum (Supplemental Table 3), including numerous human pathogens such as *Salmonella*, *Vibrio* and *Pseudomonas* spp. This systematic analysis has identified a total of 108, 871 smORFs (Supplemental Table 4), presenting an atlas of small proteins in bacteria and a tremendous increase compared to the number of known smORFs to date. We have predicted a large number of smORFs in all five classes of Proteobacteria (Figure 4A), whereas the highest number of smORFs were found among Gamma-Proteobacteria (~200 per genome).

**Figure 4.**
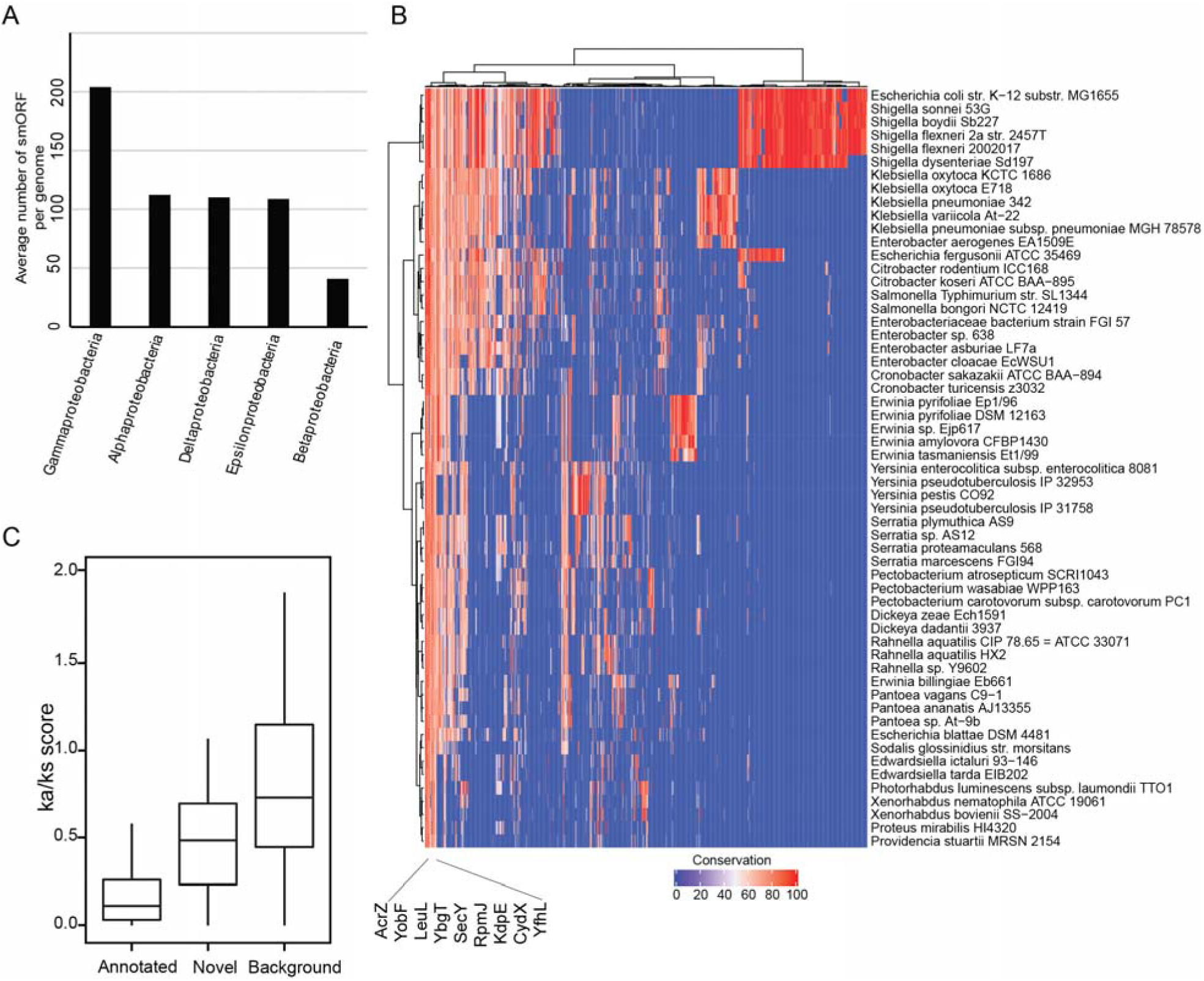
The landscape of smORFs candidates in the Proteobacteria phylum. (A) Barplot showing the average number of smORFs predicted per genome among Proteobacteria. (B) Heatmap showing the conservation profiles of smORFs in 57 representative Enterobacteria strains. Values in the heatmap represent the percentage of conservation of each smORF. (C) The distribution of Ka/Ks ratios for the known small proteins, predicted novel smORFs, and random ORF as background in *Escherichia coli* and *Salmonella*.

To better understand these novel smORFs, we have analyzed the conservation profiles for all 1235 smORFs identified in the Enterobacteriaceae family (Supplemental Table 5), blasted against a panel of 57 representative Enterobacterial species. As depicted in Figure 4B, these smORFs are separated into three major groups. The first group with highest conservation includes ribosomal associated small proteins such as RpmJ, as well as a number of known proteins such as AcrZ, YobF, LeuL, CydX and SecY. For example, CydX has been recently characterized as a conserved component of the cytochrome bd oxidase complex (Allen et al., 2014; Ramamurthi and Storz, 2014). In this group, we have identified several novel small proteins (Supplemental Table 5), whose high conservation indicates their functional importance and warrants future studies in the native bacterial strains. The second group shows somewhat intermediate conservation profiles, including small proteins identified in a range of closely-related bacterial species (Figure 4B). The third group only displays conservation patterns within one or two species, such as the strains of *Klebsiella* or *Erwinia*. A few putative uORFs were included in this group, indicating species-specific functions linked to some unique genes.

We further investigated the conservation patterns by analysing the selection pressure on codon usage (Figure 4C). For this, we have performed a Ka/Ks ratio analysis on the predicted smORFs in *Escherichia coli* and *Salmonella*, which have an average length of 25 amino acids. The result revealed that most of the novel smORFs have a substantially lower Ka/Ks ratio compared to the that of random ORFs in chromosome (Figure 4C). These predicted smORFs have a Ka/Ks ratio far below 1, indicating a significant number of nucleotide sequence changes that do not result in changes in protein sequence. In other words, the novel smORFs are under purifying selection and likely to be functional, bolstering the reliability of the sPepfinder algorithm in the prediction of novel smORFs.

## DISCUSSION

In this work, we presented a new computational approach sPepfinder that can rapidly and efficiently detect smORFs in a genome-wide manner. The sPepfinder utilizes the distinct features of bacterial small proteins, and achieves high accuracy and specificity. The application of sPepfinder in hundreds of bacterial strains has revealed the first atlas of small proteins in the Proteobacteria Phylum with over 0.1 million smORFs. With the burgeoning interest in the studies of smORFs in bacteria, the development of sPepfinder will significantly accelerate their experimental identification and characterization in a large variety of bacterial species.

sPepfinder will help identify potential smORFs in experimentally generated data, though it was initially developed for *de novo* prediction of smORFs from empty intergenic regions in genome sequences. The application of RNA-seq or ribosome profiling alone has shown promise to identify novel smORFs in model bacteria such as *E. coli* (Weaver et al., 2019). Starting from available experimental data, computational tools could more reliably predict ORFs in RNA fragments that are bound to translating ribosomes (Miravet-Verde et al., 2019; Ndah et al., 2017). A combination of sPepfinder with ribosome profiling and mass spectrometry will further enhance the accuracy of prediction, and allow the identification of functional small proteins that are highly conserved and expressed at specific physiological conditions, exemplified by our study in *Salmonella typhimurium* (Venturini et al., *in preparation*).

The sPepfinder algorithm is not without limitations. To ensure the high accuracy of prediction, we have only considered the open reading frames starting with a canonical ATG or GTG codon. The prediction will inevitable lose a small number of proteins with other non-canonical start codons (Li et al., 2012), reducing the sensitivity of prediction while maintaining a high specificity. Due to the reliance on the translation initiation score and the binding strength of RBS, our model will not take into consideration the leaderless mRNAs, though this group of mRNAs should be rare. In addition, our SVM model was trained on a collection of known small proteins in *E. coli*, therefore its performance in more distant-related organisms will need to be further evaluated, when more experimental data become available. The future inclusion of more experimentally verified small proteins will certainly improve our SVM model for a better performance and wider application in diverse bacterial species and complex bacterial communities, such as plant- and human-associated microbiome.

## MATERIAL AND METHODS

### Benchmark of small protein datasets

We have downloaded the gene table with protein only from Ecogene 3.0 database (Zhou and Rudd, 2012). In total 4323 genes were retrieved including 94 small proteins.

### The sPepfinder workflow

The intergenic sequences and with 20 bp overlap with known annotation of each genome was used as a starting point. The 20 bp upstream sequences of each smORF were extracted and the translation initiation score was calculated using Ribosome binding site calculator (Salis et al., 2009). The conservation analysis of these smORFs was performed to obtain the sequence alignment, which was further used to define the coding potential scores by RNAcode (Washietl et al., 2011). The frequency of every amino acid in small proteins and control proteins was analysed, including the three hydrophobic amino acids (I, L, V) that showed significant enrichment in known small proteins. Ribosome profiling sequencing reads were analysed by including the center weighting steps to normalize ribosome pausing sites. The *in silico* predicted smORFs are considered smORFs candidates when they show evidence of translation based on ribosome footprinting data.

### Analysis of Ribo-seq data

We overlapped our smORFs prediction result with the ribosome profiling data from *Escherichia coli* using GWIPS-viz browser (Michel et al., 2014).

### Nonsynonymous and synonymous substitutions (Ka/Ks ratio)

The nucleotides and peptide sequences of small peptides in *Escherichia coli* and *Salmonella* Typhimurium were analysed using the KaKs calculator software (Zhang et al., 2006) to examine nonsynonymous (Ka) and synonymous (Ks) substitutions and the resulting and Ka/Ks values, using the approximate method ‘NG’.

## ACKNOWLEDGEMENT

We thank Konrad Förstner for his initial input and suggestions, and Jörg Vogel for critical reading and comments on the manuscript.

## AUTHOR CONTRIBUTIONS

L.L., Y.C. conceived and supervised this project. L.L. performed the data analyses. L.L., Y.C. interpreted the data and wrote the manuscript.

## CONFLICT OF INTEREST

The authors declare no competing financial interests.

## SUPPLEMENTAL TABLES

**Table S1.** The training datasets used for sPepfinder. The SVM model was trained using a positive dataset of 123 known small proteins, and a negative dataset including 136 short ORFs predicted in rRNA and tRNA regions.

**Table S2.** The list of 124 smORFs predicted in *E. coli* by sPepfinder.

**Table S3.** The list of all the 549 proteobacterial genomes used for prediction.

**Table S4.** The complete list of all the predicted smORFs in Proteobacteria.

**Table S5.** The list of the conserved smORFs in 57 Enterobacteria.

## Notes

### Competing Interest Statement

The authors have declared no competing interest.

